# The La Crosse virus class II fusion glycoprotein *ij* loop contributes to infectivity and cholesterol-dependent entry

**DOI:** 10.1101/2023.02.22.529620

**Authors:** Sara A. Thannickal, Sophie N. Spector, Kenneth A. Stapleford

## Abstract

Arthropod-borne viruses (arboviruses) are an emerging and evolving global public health threat with little to no antiviral treatments. La Crosse virus (LACV) from the *Bunyavirales* order is responsible for pediatric encephalitis cases in the United States, yet little is known about the infectivity of LACV. Given the structural similarities between class II fusion glycoproteins of LACV and chikungunya virus (CHIKV), an alphavirus from the *Togaviridae* family, we hypothesized that LACV would share similar entry mechanisms to CHIKV. To test this hypothesis, we performed cholesterol-depletion and repletion assays and used cholesterol modulating compounds to study LACV entry and replication. We found that LACV entry was cholesterol-dependent while replication was less affected by cholesterol manipulation. In addition, we generated single point mutants in the LACV *ij* loop that corresponded to known CHIKV residues important for virus entry. We found that a conserved histidine and alanine residue in the Gc *ij* loop impaired virus infectivity and attenuate LACV *in vitro* and *in vivo*. Finally, we took an evolution-based approach to explore how the LACV glycoprotein evolution in mosquitoes and mice. We found multiple variants that cluster in the Gc glycoprotein head domain, supporting the Gc glycoprotein as a target for LACV adaptation. Together, these results begin to characterize the mechanisms of LACV infectivity and how the LACV glycoprotein contributes to infectivity and pathogenesis.

**Importance:** Vector-borne arboviruses are significant health threats leading to devastating disease worldwide. This emergence and the fact that there are little to no vaccines or antivirals targeting these viruses highlights the need to study how arboviruses replicate at the molecular level. One potential antiviral target is the class II fusion glycoprotein. Alphaviruses, flaviviruses, and bunyaviruses encode a class II fusion glycoprotein that contain strong structural similarities in the tip of domain II. Here we show that the bunyavirus La Crosse virus uses similar mechanisms to entry as the alphavirus chikungunya virus and residues in the *ij* loop are important for virus infectivity. These studies show that genetically diverse viruses use similar mechanisms through concerned structure domains, suggesting these may be a target for broad-spectrum antivirals to multiple arbovirus families.

## Introduction

Arthropod-borne viruses (arboviruses) are significant human pathogens with epidemic potential (1–3). Arboviruses must infect and replicate within humans and insects, yet the molecular mechanisms these viruses use for infectivity and replication are not completely understood. Arboviruses include a long list of human pathogens that span multiple virus families. These families include the positive-strand RNA alphaviruses and flaviviruses as well as the segmented negative-strand RNA bunyaviruses. While our knowledge of how alphaviruses and flaviviruses infect and replicate in mammals and insects has expanded in response to large outbreaks, we know little about how bunyaviruses infect these very disparate hosts.

The *Bunyavirales* order includes a number of significant human pathogens such as Rift Valley fever virus, La Crosse virus, Oropouche virus, and Crimean Congo hemorrhagic fever virus (4, 5). La Crosse virus (LACV) is a member of the orthobunyavirus genus and is an enveloped segmented negative-strand RNA virus that contains a tripartite genome consisting of S, M, and L segments (6, 7). The S segment encodes for the nucleoprotein (N) and NSs protein (4, 8), the M segment encodes for the viral glycoproteins Gn and Gc (7, 9) and nonstructural protein NSm, and the L segment encodes for the viral RNA dependent RNA polymerase (4). LACV is transmitted by the *Aedes* species of mosquito (10, 11) and can cause severe encephalitis, as it is the most common cause of pediatric neuroinvasive arboviral disease in children in the United States (12–15).

However, while our knowledge of bunyavirus entry is limited, we can use what we know from alphaviruses and flaviviruses to infer how bunyaviruses establish infections. Alphaviruses, flaviviruses, and bunyaviruses encode class II fusion glycoproteins required for assembly, entry, and virion stability (16–18). While the amino acid sequences between viral families show minimal sequence homology, the class II glycoproteins of these three genetically distant viral families share striking structural homology in domain II, which includes the fusion loop required for membrane fusion (18) (**Fig. 1**). In addition to the fusion loop, each class II fusion protein contains an *ij* and *bc* loop flanking the fusion tip. Extensive work with alpha- and flaviviruses has shown that the *ij* loop is critical for virus infectivity (19), cholesterol-dependent entry (20) (21–24), and transmission (25, 26). In addition, the *ij* loop and β-strand c have been shown to be key sites of adaptive evolution leading to increased transmission in mosquitoes for both loops, while β-strand c has been associated with enhanced virulence (27, 28). Given the structural similarities between these viral proteins we hypothesize that the bunyavirus *ij* loop will also play key roles in virus replication.

**Figure 1:**
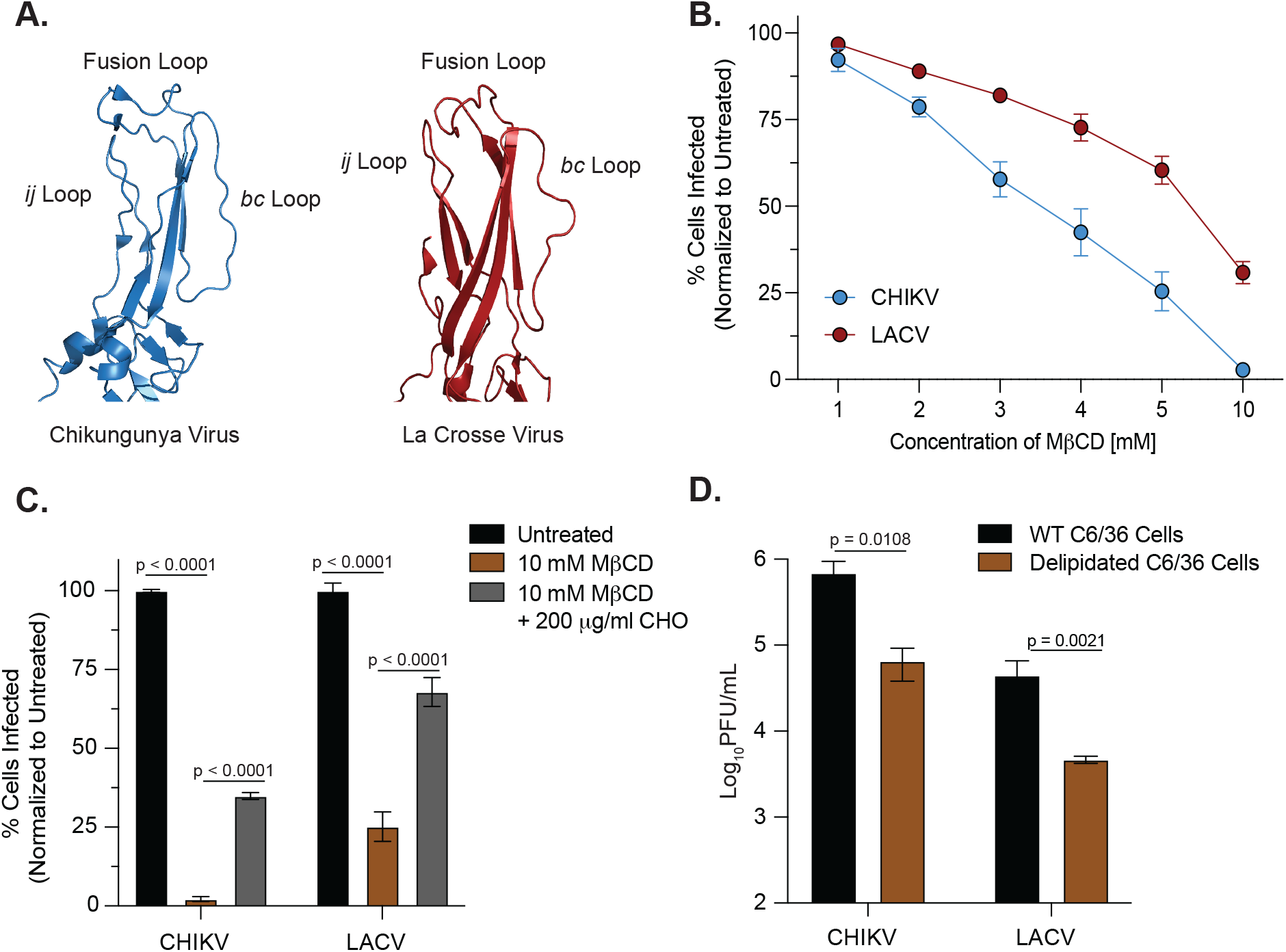
La Crosse virus entry is cholesterol-dependent. (A) Crystal structures of domain II of the chikungunya virus (CHIKV) E1 (PDB: 3N42) and La Crosse virus (LACV) Gc (PDB: 7A57) fusion glycoproteins with the *ij* loop, fusion loop and *bc* loop labeled. (B) BHK-21 cells were pretreated with methy-β-cyclodextrin (MβCD) and cholesterol-depleted cells were infected with CHIKV-ZsGreen or LACV (Human/78) at an MOI of 1 before adding 20 mM NH_4_Cl. Cells were fixed at 24 hours post-infection. LACV infections were stained with primary antisera and a goat anti-rabbit IgG-Alexa488 secondary antibody, and both infections were stained with DAPI. The number of infected cells were quantified using a CX7 high-content microscope. Data are normalized to an untreated control and represent three independent trials with internal duplicates. (C) BHK-21 cells were pretreated with 10 mM MβCD and were incubated with either DMEM or 200 μg/mL water-soluble cholesterol. Cholesterol-depleted, complemented, or untreated cells were infected with CHIKV-ZsGreen or LACV (Human/78) at an MOI of 1 for 1 hour before adding 20 mM NH_4_Cl. Data represent two independent experiments with internal triplicates. A Two-Way Anova test was performed and p-values are shown. (D) Delipidated C6/36 insect cells were infected with CHIKV (MOI = 0.01) or LACV (MOI = 0.1). After a PBS wash and adding back serum media, viral supernatants were harvested 24 hours post-infection and used to quantify infectious particles by plaque assay. Data represent two independent experiments with internal triplicates. Mann-Whitney tests were performed and p-values are shown. The average and standard error of the mean (SEM) are shown for all data.

In this study, we use LACV as a model orthobunyavirus to investigate entry and the role of the *ij* loop in the bunyavirus life cycle. We show that LACV entry, similar to alphaviruses, is cholesterol-dependent. However, while alphavirus replication seems to rely on cholesterol, LACV replication is less affected. Using an infectious clone, we generated point mutants in the LACV *ij* loop and found that specific residues contributed to overall infectivity as well as cholesterol-dependent entry. Importantly, the *ij* loop variants were attenuated *in vitro* in BHK-21 cells and the variant H1193A was mildly attenuated in a newborn mouse model of infection.

Finally, given that the alphavirus glycoprotein has been found as fundamental for adaptive evolution, we addressed the evolution of the LACV glycoproteins. In mice and mosquitoes, we found a number of mutations in the Gc head domain, suggesting it may be important for host adaptation. Taken together, our data show that bunyaviruses use similar mechanisms, domains, and amino acids for virus entry as alphaviruses. Moreover, we have identified the Gc head domain as a hotspot for virus adaptation *in vivo*. These studies and future work dissecting the bunyavirus glycoprotein are critical to our global understanding of class II glycoprotein function and viral pathogenesis.

## Results

### Host cholesterol contributes to La Crosse virus entry

Alphaviruses, flaviviruses and bunyaviruses all encode a class II fusion glycoprotein that play a vital role in mediating viral attachment and membrane fusion (7, 16, 17). However, while our understanding of alphavirus and flavivirus glycoprotein biology have grown significantly, our understanding of vector-borne bunyavirus glycoprotein function is less well understood. To begin to understand how the bunyavirus glycoprotein functions, we first explored the structural similarities between the class II fusion glycoprotein of the orthobunyavirus La Crosse virus (LACV) and the alphavirus chikungunya virus (CHIKV) (**Fig. 1A**). Given the structural similarities in the tip of domain II between these two glycoproteins, particularly the fusion loop, *ij* loop and *bc* loop, we hypothesized that LACV would use a cholesterol-dependent mechanism of entry similar to what has been shown for CHIKV. To investigate the cholesterol-dependent entry of LACV, we depleted BHK-21 cells of cholesterol using increasing concentrations of methyl-β-cyclodextrin (MβCD), and then infected cells with either wild-type CHIKV-ZsGreen or LACV (Human/78 strain) (**Fig. 1B**). We found that both CHIKV and LACV were sensitive to cholesterol depletion, with CHIKV being more sensitive than LACV. Notably, we were able to restore CHIKV or LACV infection when we added back cholesterol to the BHK-21 cells, yet not to the same extent as our untreated controls (**Fig. 1C**). Finally, since both CHIKV and LACV infect *Aedes* (*Ae*.) mosquitoes, we wanted to test their cholesterol-dependent entry in C6/36 *Ae. albopictus* cells. We grew C6/36 cells in the present or absence of delipidated serum and tested whether delipidation affected viral entry (**Fig. 1D**). We found both viruses sensitive to delipidation, indicating that LACV infection is cholesterol-dependent in both mammalian and mosquito cells. These data show that both CHIKV and LACV utilize cholesterol for entry, as depleting levels of cholesterol and lipids decrease the infectivity of each wild type virus.

### Cholesterol modulating compounds have little impact on LACV replication

As an alternative approach to understanding the role of cholesterol in the LACV life cycle, we used the cholesterol-modulating compounds atorvastatin, lovastatin and imipramine. Atorvastatin and lovastatin are FDA-approved HMG-CoA reductase inhibitors that decrease cholesterol levels in the liver while also stimulating LDL receptors expression on hepatic cells (29, 30). Imipramine is an FDA-approved anti-depressant that disrupts cholesterol trafficking within the cell. As these compounds have been shown to inhibit the entry and replication of CHIKV and other arboviruses (31–34), we hypothesized they may also alter LACV infection as well. We first tested the cytotoxicity of each compound using an MTT assay and observed limited cytotoxicity in BHK-21 cells until 25 and 50 μM for all compounds except our DMSO control (**Fig. 2A**). We then pretreated BHK-21 cells with each compound for two hours and infected the cells with wild-type CHIKV or LACV and added media containing each compound for 24 hours. We found that CHIKV and LACV were sensitive to imipramine in BHK-21 cells, however at higher concentrations the decrease in infected cells may be the result of cytotoxicity from the compound alone (**Fig. 2B**), while atorvastatin and lovastatin had minimal effect on the infectivity of each virus (**Fig. 2C** and **2D**). These results suggest that although both LACV and CHIKV are dependent on cholesterol for entry, they may use cholesterol differently for replication.

**Figure 2:**
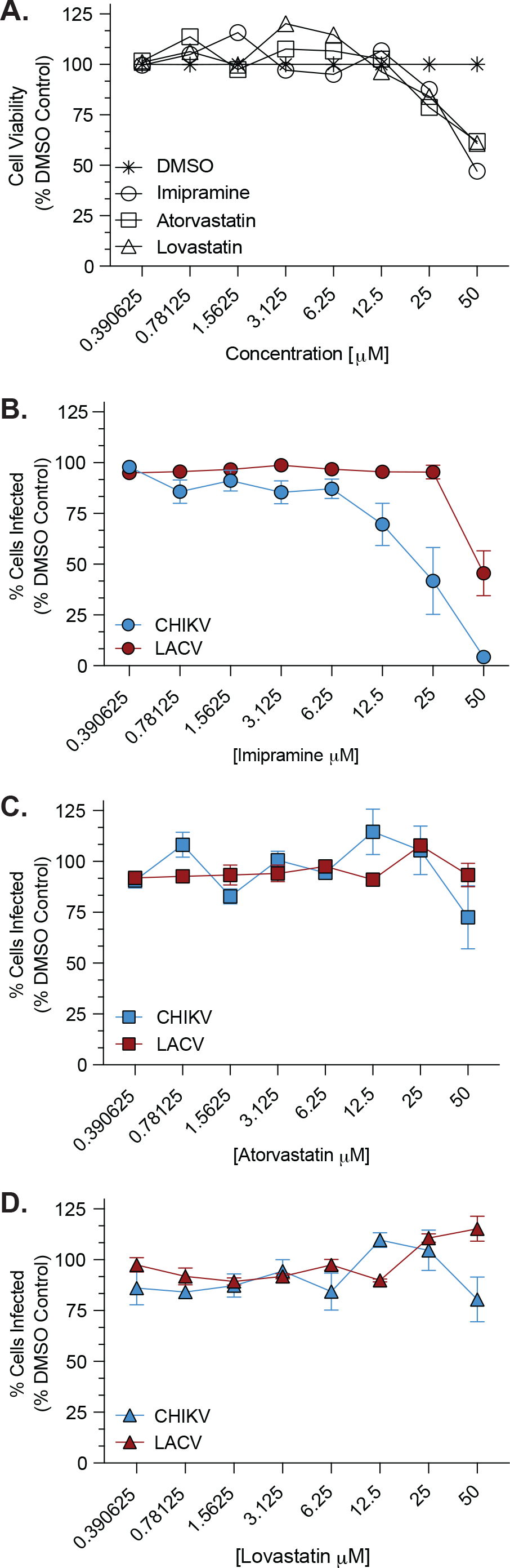
LACV shows selective sensitivity to cholesterol-modulating compounds. (A) An MTT assay was used to determine cytotoxicity from Imipramine, Atorvastatin and Lovastatin in BHK-21 cells at increasing concentrations at 36 hours post-treatment. BHK-21 cells were pre-treated with Imipramine (B), Atorvastatin (C), or Lovastatin (D) at increasing concentrations and then infected with CHIKV-ZsGreen or LACV (Human/78) at an MOI of 0.1 for 1 hour. Cells were then washed with PBS and incubated in media with the same drug concentration for an additional 24 hours before being fixed and stained for analysis using a CX7 high-content microscope. Data are normalized to a DMSO control and represent at least two independent experiments with internal duplicates. The average and standard error of the mean (SEM) are shown for all data.

### LACV Gc *ij* loop is important for LACV replication *in vitro* and *in vivo*

Domain II of the class II fusion glycoproteins of alpha-, flavi-, and bunyaviruses contains the fusion loop (*cd* loop), the *ij* loop, and the *bc* loop. While the fusion loop of La Crosse has been studied extensively (7, 35–37), how the other loops function in virus entry has not been explored. In particular, the *ij* loop of alphaviruses and flaviviruses has been shown to be critical for virus assembly, entry, and transmission (19, 24, 26). In addition, we have shown that a conserved alanine residue in the Zika virus and yellow fever virus *ij* loop are critical for virion assembly and envelope protein accumulation (38). Therefore, we hypothesized that the orthobunyavirus *ij* loop would also play important roles in assembly and infectivity. To address this hypothesis, we first looked at the conservation of the *ij* loop between orthobunyaviruses (**Fig. 3A**). Interestingly, several orthobunyaviruses, including LACV have an alanine at the tip of the *ij* loop, similar to alpha- and flaviviruses. In addition, there is a conserved histidine and arginine flanking the *ij* loop tip, suggesting these may be important for function. Given this conservation between viruses and viral families, we generated point mutations in the *ij* loop of the Gc glycoprotein of LACV (LACV/78/NC-cl strain), which we hypothesized would be vital for infection *in vitro* and *in vivo* (**Fig. 3A and 3B**). To investigate the role of these residues in the LACV life cycle, we first performed growth curves in BHK-21 cells and found that all three *ij* loop point mutants were attenuated at 16 and 24 hours post-infection indicating that the *ij* loop is important in the LACV life cycle (**Fig. 3C**). We then asked whether each mutant is attenuated *in vivo* by infecting 7-8 day old C57BL/6 mice with LACV WT or each *ij* loop mutant and quantifying viral titers in the brain 3 days post-infection (**Fig. 3D**). We found that within each trial, each mutant was slightly attenuated compared to wild-type, but the H1193A mutant was consistently attenuated in mice compared to the other mutants. These results indicate that the *ij* loop is important for LACV infection *in vitro* and *in vivo*, similar to what was found for LACV fusion loop variants (36, 37).

**Figure 3:**
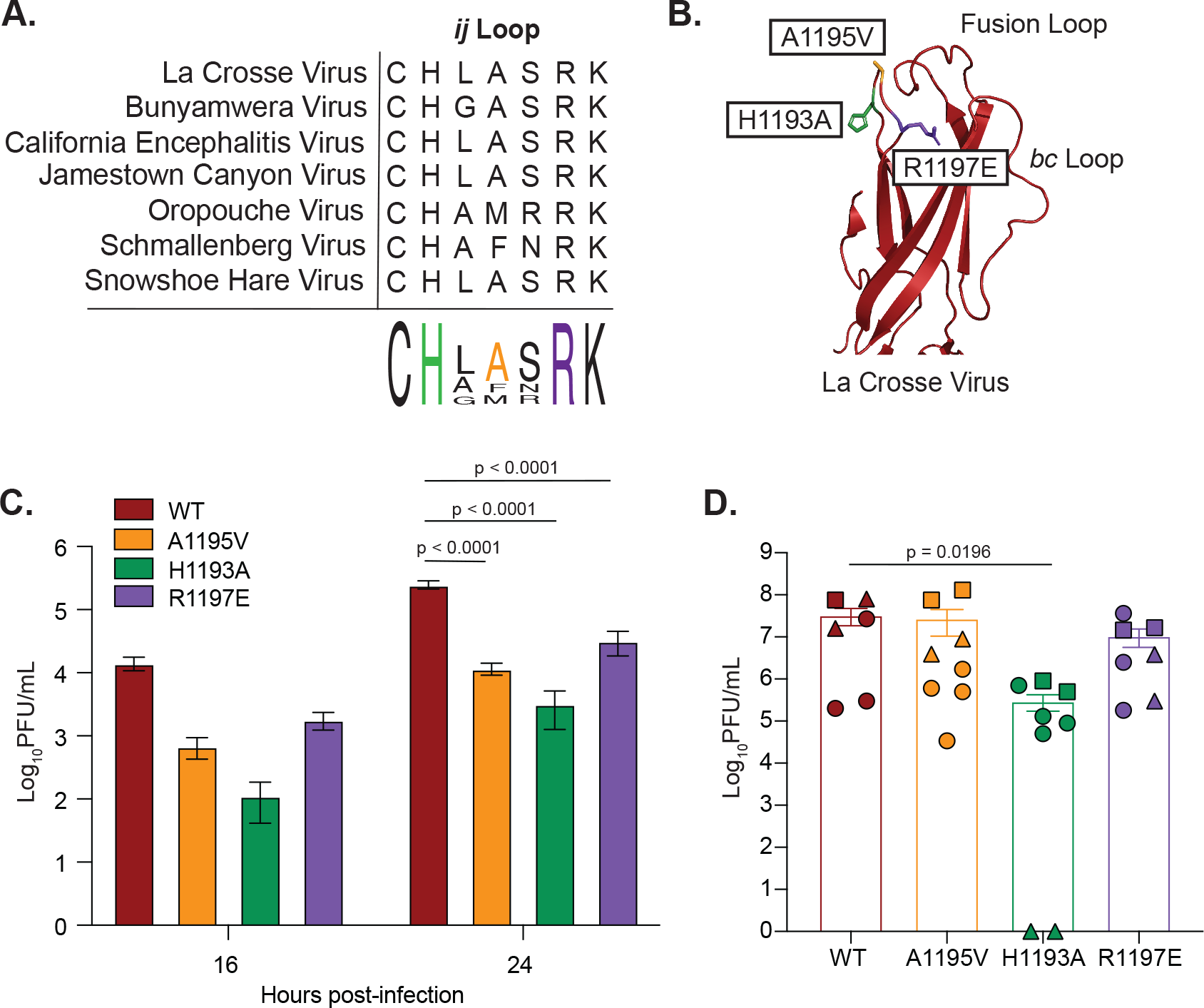
The Orthobunyavirus Gc *ij* loop is highly conserved and contributes to LACV infection. (A) Protein alignment of the *ij* loop of different orthobunyaviruses. (B) Crystal structure of the LACV Gc domain II showing individual *ij* loop mutations (PDB: 7A57). (C) BHK-21 cells were infected with wild-type LACV (78/NC-cl) or each variant at an MOI of 0.1 for 1 hour. Culture supernatants were collected at the indicated time points and viral titers were quantified by plaque assay. Data represent at least two independent experiments with internal duplicates. A Two-Way Anova test was performed and p-values are shown. (D) 7-8 day old C57BL/6J mice were infected subcutaneously with 50 PFU of either LACV WT or each *ij* loop mutant. Mice were euthanized 3 days post inoculation, the brain was harvested and used for a plaque assay to quantify infectious particles. Shapes correspond to a different cage of pups that were infected, and represent 3 independent experiments with at least n=6 mice for each condition. A Mann-Whitney test was performed and data with statistical significance are labeled with their corresponding p-value. The average and standard error of the mean (SEM) are shown for all data.

### The LACV Gc *ij* loop contributes to infectivity and cholesterol-dependent entry

The alphavirus *ij* loop has been shown to contribute to infectivity, cholesterol-dependent entry, and transmission. Notably, the emergence of an A226V variant in the *ij* loop of the E1 glycoprotein of CHIKV contributed to the 2005 La Reunion epidemic and has been shown to contribute to cholesterol-dependent entry (23, 26, 39). Given these findings, we asked whether the LACV *ij* loop behaved similarly. To begin, we first addressed virus infectivity by infecting BHK-21 cells with either LACV WT or each *ij* loop mutant at an MOI of 1 for 1 hour and then added 20 mM ammonium chloride to each cell for a 24-hour incubation to block virus spread (**Fig. 4A and 4B**). After fixing and staining the cells, we found the H1193A and R1197E variants increased infectivity compared to WT, while the A1195V variant lead to reduced infectivity in BHK-21 cells. Noting their differences in infectivity, we then wanted to test the *ij* loop’s role in cholesterol-dependent entry. We depleted BHK-21 cells of cholesterol using MβCD as before and infected cells with either LACV WT or each *ij* loop mutant (**Fig. 4C and 4D**). We found that the while the infectivity of the mutants varied (**Fig. 4A**), variants H1193A and R1197E showed a more extreme decay in infectivity after treatment compared with wild-type, while the A1195V did not seem to reduce infection any further than the untreated control (**Fig. 4D**). Our findings show that changes to the *ij* loop led to variation in infectivity and cholesterol dependency in BHK-21 cells *in vitro* compared to WT, reiterating the importance of the *ij* loop for LACV infection.

**Figure 4:**
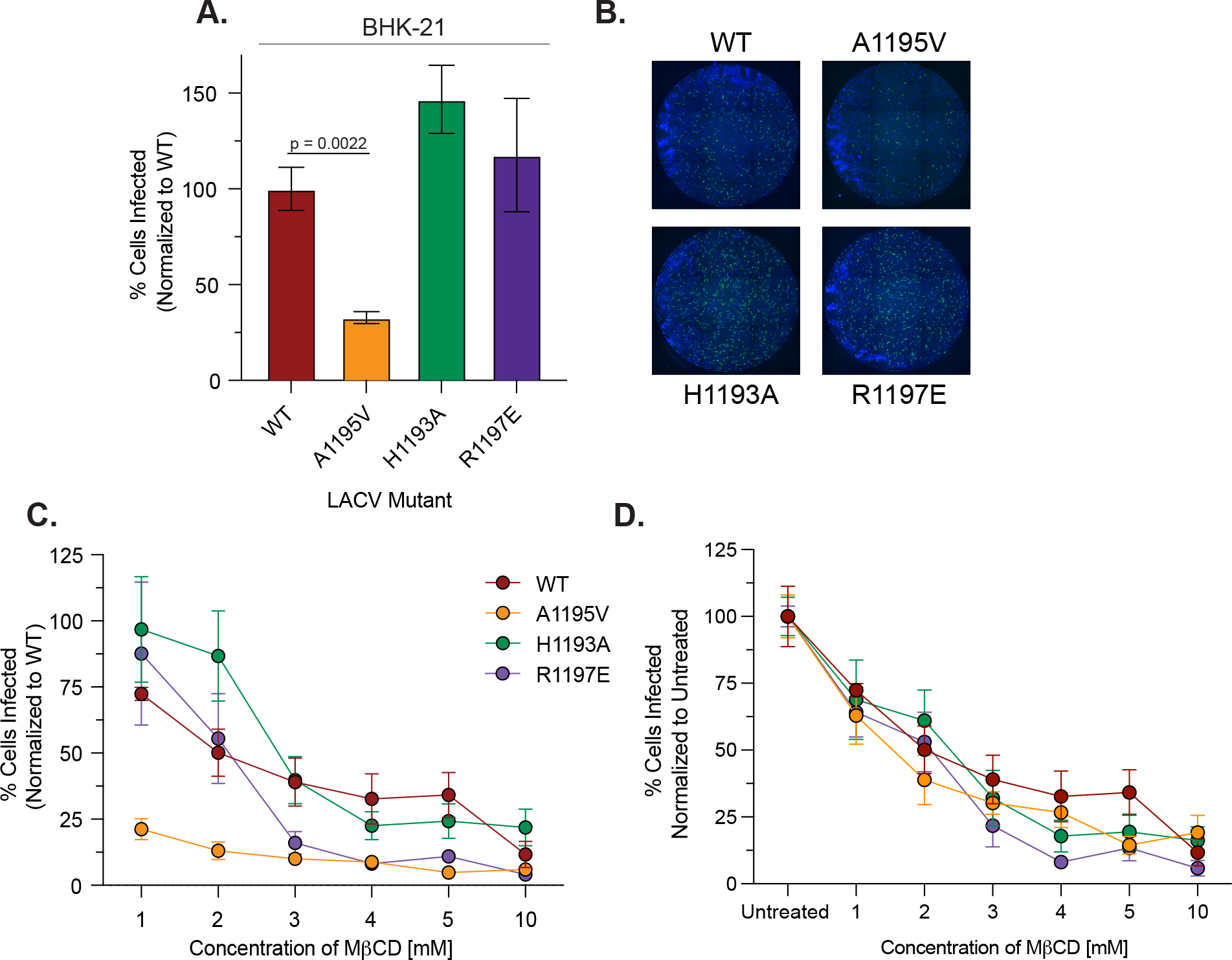
LACV Gc *ij* loop variants contribute to LACV infectivity and cholesterol-dependent entry. (A) BHK-21 cells were infected with either WT LACV (78/NC-cl) or each *ij* loop mutant at an MOI of 1 for 1 hour. After incubation, 20 mM NH_4_Cl was added and the cells incubated for 24 hours before being fixed and stained for analysis. Data are normalized to WT LACV and represent three independent trials with internal duplicates. A Mann-Whitney test was performed and data with statistical significance are labeled with their corresponding p-value. (B) Representative images from a CX7 high-content microscope with the nuclei of BHK-21 cells stained blue with DAPI and LACV infected cells green from the antisera primary antibody and a goat anti-rabbit IgG-Alexa488 secondary antibody. (C and D) BHK-21 cells were pretreated with methy-β-cyclodextrin (MβCD) and cholesterol-depleted cells were infected with LACV WT or each *ij* loop mutant at an MOI of 1. Cells were then treated with 20 mM NH_4_Cl for 24 hours and were then fixed, stained and quantified as before. Data are normalized to an untreated WT or untreated control within each mutant, and represent three independent experiments with internal duplicates. The average and standard error of the mean (SEM) are shown for all data.

### *In vivo* evolution of LACV highlights the Gc head domain as potential hotspot for adaptation

The arbovirus glycoproteins are key determinants of virus adaptation (25, 40). Experiments in the lab and in nature have shown that the viral glycoproteins can adapt *in vivo* (27, 38, 39, 41), providing key snapshots into glycoprotein function. Therefore, we hypothesized that *in vivo* LACV infections may shed light on how LACV may evolve. Specifically, mosquitoes play a fundamental role in the evolution of emerging arboviruses, as the interaction with the host vector can cause various mutations to emerge in nature and change the biology of these viruses. LACV has been found to infect *Aedes* (*Ae*.) species of mosquitoes, including *Ae. albopictus*, through surveillance of different areas in the United States (42). To address the *in vivo* evolution of LACV (Human/78 strain) we infected *Ae. aegypti* (Mexico) and *Ae. albopictus* (New York) via an artificial blood meal. As a control for infection, we included CHIKV and as its primary vectors are *Ae. aegypti* and *Ae. albopictus*. At 7 days post-infection, we harvested individual mosquitoes and separated bodies (infection) from legs and wings (dissemination), then determined infectious titers via plaque assay (**Fig. 5A and 5B**). While CHIKV infected and disseminated in both species, LACV preferentially infected *Ae. albopictus*, yet poorly disseminated in both species. We then extracted the RNA from the bodies of the LACV infected *Ae*. mosquitoes and Sanger sequenced the full LACV genome (**Fig. 5C**). Interestingly, we found a few synonymous and non-synonymous mutation in the S and L segments of individual mosquitoes (**Fig. 5C**). However, in the M segment, where the glycoprotein is encoded, we found a number of mutations including one (R618W/L) that was found in multiple mosquitoes (**Fig. 5C and 6A**). Finally. in addition to mosquitoes, we infected 5-10 day old mice with LACV (Human/78 strain) to understand LACV evolution in mammals. In mice, we found no nonsynonymous or synonymous mutations in the S or L segment. However, in the M segment we found a number of mutations including several in the Gc head domain (R618W, D619G, A621V, and Q795R) present in nearly all infected mice (**Fig. 6A**). When we mapped these variants onto the crystal structure of the Gc head domain we found these mutants to cluster at the interface of the head domain, which is important for contacts between Gc monomers of the trimer (43) (**Fig. 6B, C, and D**). Together, these results indicate that the M segment, and specifically the Gc head domain, may be a potential hotspot for viral evolution, as the frequent mutations found suggest the importance of these residues for infection and transmission in nature.

**Figure 5:**
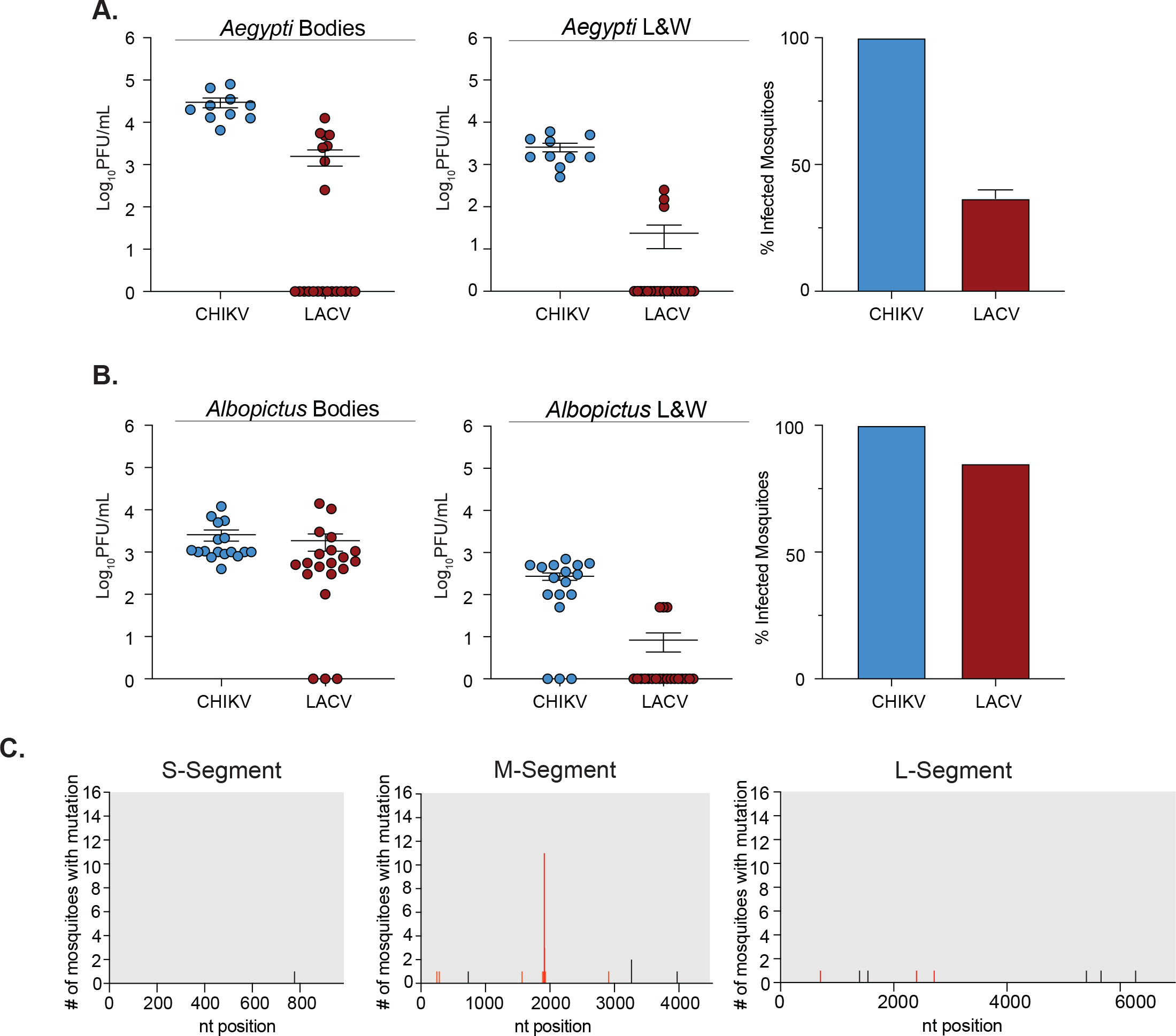
LACV mosquito infections and evolution. *Aedes (Ae) aegypti* (A) or *Ae. albopictus* (B) mosquitoes were infected with 10^6^ PFU/mL of CHIKV or LACV (Human/78) via an artificial blood meal. Mosquitoes were harvested and dissected 7 days post-infection, and infectious titers were determined by plaque assay in the bodies or legs and wings. Perfect of infected mosquitoes are shown on the right. Data represents at least one independent experiment with at least n=10 mosquitoes for each condition. The average and standard error of the mean (SEM) are shown for all data. (C) The full LACV genome of individual mosquitoes was Sanger sequenced and graphs represent the frequency of mutations found within each LACV genome segment. Black tick marks indicate a synonymous mutation, and red tick marks indicate a non-synonymous mutation with its position labeled.

**Figure 6:**
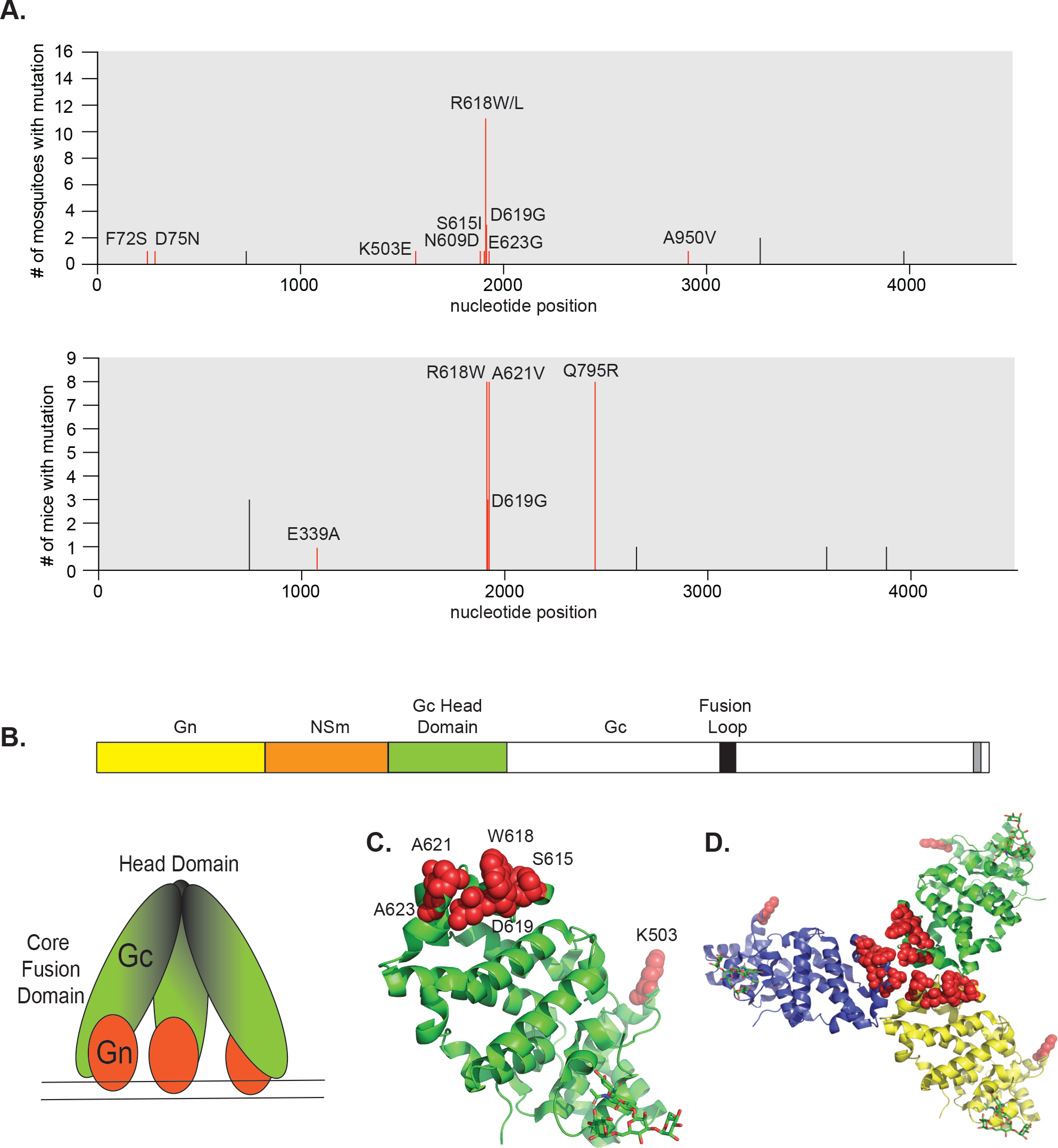
The LACV Gc head domain is a potential hotspot for evolution. (A) Frequency of mutations found in the LACV M segment by Sanger sequencing from mosquitoes and mice. Black tick marks indicate a synonymous mutation, and red tick marks indicate a non-synonymous mutation with its position on the glycoprotein labeled. (B) A schematic showing the LACV M segment, as well as a depiction of the Gc trimer head domain and Gn spike. Crystal structure of the Gc head domain with red spheres indicating the mutations found from the *in vivo* Sanger sequencing results as a monomer (C) or trimer (D) (PDB: 6H3W).

## Discussion

Arboviruses are genetically diverse pathogens that can cause devastating disease worldwide. There are limited antivirals or vaccines that target these viral threats, highlighting the need to study arbovirus biology at the molecular level. The majority of arboviruses fall into three major viral families, the *Togaviridae* (alphavirus genus), *Flaviviridae* (flavivirus genus), and the *Bunyavirales* order. While extensive work has been done to study alphavirus and flavivirus biology, there is still much unknown about bunyavirus mechanisms of entry, pathogenesis and evolution, despite high epidemic potential.

La Crosse virus (LACV) is an emerging neuroinvasive arbovirus that has been a major cause of pediatric encephalitis throughout the United States, with persistent high-risk clusters of cases in the Midwest and Appalachian regions (12). Here, we used the structural similarities of the CHIKV E1 and the LACV Gc class II fusion glycoproteins to hypothesize the importance of the *ij* loop for cholesterol-dependent entry and infectivity *in vitro* and *in vivo*. Like alphaviruses, we found that La Crosse virus entry was cholesterol-dependent in both mammalian and insect cell lines. While CHIKV shows to be more sensitive, both CHIKV and LACV show a decrease in infected cells with depleting levels of cholesterol and lipids, signifying the importance of cholesterol for LACV entry. However, in the presence of cholesterol-modulating compounds, both CHIKV and LACV seem to be less sensitive to treatment, as only Imipramine decreased infected cells in BHK-21 cells, though the cytotoxicity of the drug at high concentrations may play a factor in these results. These results may be due to differences in host and cell type as previous work with impramine and statins were completed in human cells (33, 34). These results show that while LACV entry is cholesterol-dependent similar to CHIKV, replication in LACV is less affected by cholesterol manipulating treatments. Characterizing LACV’s cholesterol dependence is crucial, as a re-emergent CHIKV strain from the Indian Ocean islands outbreak shifted its dependency on cholesterol and enhanced transmissibility in *Ae. albopictus* (26). The increased transmission of CHIKV during this epidemic highlights the importance of monitoring LACV and other arboviruses for mechanisms of entry and replication that could be enhanced due to single or multiple point mutations.

The alphavirus E1 A226 residue resides on the *ij* loop, which has been characterized extensively as important for infectivity (19, 22), cholesterol-dependence (24) and transmission (25, 26). The *ij* loop of orthobunyaviruses is highly conserved and contains a conserved histidine and arginine, as well as an analogous alanine to the alphavirus A226 residue. We found that all three *ij* loop variants were attenuated in BHK-21 cells with the A1195V mutant having reduced infectivity, potentially explaining its attenuated growth *in vitro*. It is interesting that the H1193A and R1197E variants had no impact on infectivity, yet still showed reduced growth *in vitro*, suggesting they may play other roles in the viral life cycle. However, in neonatal mice, only the H1193A is attenuated, with one trial having no infectious particles found in the brain. A similar histidine in the alphavirus *ij* loop has been shown to play important roles in Semliki Forest virus fusion (19), suggesting a similar role in LACV. Future studies are underway to continue to further dissect the mechanisms of how the ij loop functions *in vitro* and *in vivo*.

Although the primary vector of La Crosse virus is the *Ae. triseriatus* mosquito, LACV has been found to infect other *Aedes* species, including *Ae. albopictus*, *Ae. japonicus*, and *Ae. aeygpti* (11, 44). Here, we demonstrate that LACV (Human/78 strain) can infect and disseminate in both *Ae. aegypti* and *Ae. albopictus* mosquitoes, however not to the same extent as CHIKV. Importantly, given the need for *in vivo* models to study arboviral evolution, we found that sequencing LACV from infected mice and mosquitoes identified multiple point mutations in the M segment, specifically in the Gc head domain. One mutation in particular at residue R618 has been found previously during mosquito passage, indicting this residue could be important for vector infections (45). There have been few studies highlighting the importance of the Gc head domain, yet it has been shown to be highly immunogenic and a potential target as a vaccine against another orthobunyavirus Schmallenberg virus (46). Studying the LACV Gc head domain as a potential hotspot for evolution is essential as these prominent residues in our studies reveal how LACV can evolve and emerge in different vector hosts.

Together, our studies highlight the importance of the LACV Gc *ij* loop in the LACV life cycle *in vitro* and *in vivo*. The highly conserved residues found in the orthobunyavirus Gc *ij* loop are critical for entry, infectivity and evolution of this emerging virus. Further studies are underway to evaluate these and other Gc residues and their importance on LACV pathogenesis and transmission *in vitro* and *in vivo*.

## Acknowledgements

We thank all members of the Stapleford Lab for helpful comments on this project. We thank Dr. Meike Dittmann at the New York University Grossman School of Medicine (NYUGSoM) for use of the CX7 high-content microscope and Drs. Ludovic Desvignes and Dominick Papandrea for their support with the NYU ABSL3 high-containment laboratory. This work was supported by funding from the NYUGSoM Start up and NIAID/NIH R01 AI162774-01A1.

## Materials and Methods

### Cell lines

BHK-21 cells (ATCC CCL-10) were grown in Dulbecco’s Modified Eagle Medium (DMEM; Corning) with 10% fetal bovine serum (FBS, Atlanta Biologics), 1% nonessential amino acids (NEAA, Corning) and 1% HEPES (Invitrogen). BHK-BSR/T7 cells were a gift from Dr. Steven Whitehead (National Institutes of Health (NIH)) and maintained in the same media as the BHK-21 cells above with 1 mg/mL gentamicin (Gibco) added every other passage (47). Vero cells (ATCC CCL-81) were grown in DMEM with 10% newborn calf serum (NBCS, Sigma). Mammalian cells were maintained at 37°C with 5% CO_2_. C6/36 *Ae. albopictus* cells (ATCC CRL-1660) were maintained in L-15 Lebovitz Medium with 10% FBS, 1% NEAA, 1% tryptose phosphate broth at 28°C with 5% CO_2_. All cell lines were confirmed to be mycoplasma free.

### Viruses

La Crosse virus (LACV) strain Human/78 was obtained through the NIH Biodefense and Emerging Infections Research Resources Repository, NIAID, NIH (NR-540). The LACV strain LACV/78/NC-cl infectious clone was a gift from Dr. Steven Whitehead (NIH) (47). LACV Gc *ij* loop variants were generated by overlapping Phusion PCR using the primers in **Table 1**. In brief, a ~2.4 kb overlapping PCR product containing 5’ EcoRI and 3’ NotI restriction sites was generated and subcloned into the same restriction sites of the wild-type plasmid. All plasmids were sequenced in full at Plasmidsaurus. To generate infectious virus, BHK BSR/T7 cells were transfected with 2 μg of each T7-driven plasmid encoding S, M, and L segments using TransIT-LT1 reagent (Mirus) following the manufacturer’s instructions (47). 24 hours post-transfection, the media was replaced with fresh media and cells incubated at 37°C for 4 days. To generate a working stock of all viruses, LACV was amplified on Vero CCL-81 cells, the supernatant clarified by centrifugation, and the virus was aliquoted and stored at −80°C. Infectious viral titers were quantified by plaque assay as described below. The entire coding region of the LACV genome (S, M, and L segments) was Sanger sequenced at Genewiz. Sanger sequences were used to generate reference genomes for variant detection.

**Table 1:**
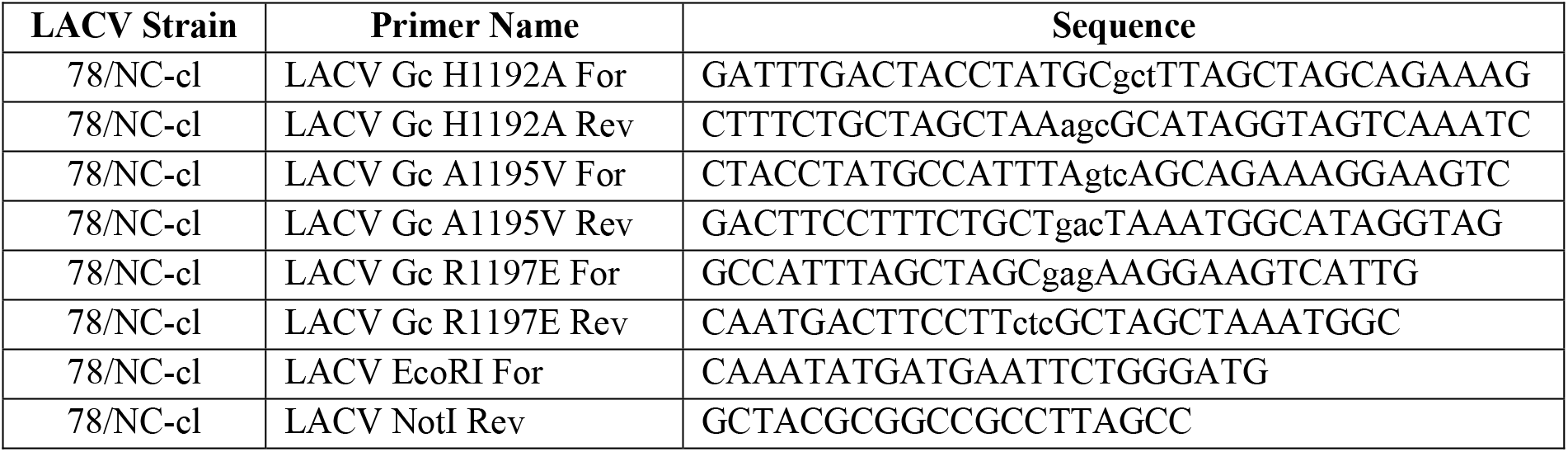
Site directed mutagenesis primers used in this study. (lowercase = mutant codon)

The chikungunya virus (CHIKV) strain 06-049 (accession #AM258994) infectious clone encoding a ZsGreen reporter was previously described (28). 10 μg of the infectious clone plasmid was linearized overnight with NotI, purified by phenol:chloroform extraction and ethanol precipitation, and resuspended in nuclease free water. CHIKV-ZsGreen RNA was *in vitro* transcribed using the mMessage mMachine SP6 kit (Invitrogen) following the manufacturer’s instructions. RNA was purified by phenol:chloroform extraction and ethanol precipitation, diluted to 1 μg/μl, aliquoted and stored at −80°C. To generate infectious virus, BHK-21 cells (10^7^/mL) were electroporated with 10 μg of *in vitro* transcribed RNA by 1 pulse of 1,200 V, 25 Ω, and infinite resistance. Electroporated cells were added to a T25 flask in BHK-21 media (DMEM, 10% FBS, 1% NEAA and 1% HEPES) and incubated at 37°C for 72 hours.

After incubation, the supernatant was centrifuged at 1,200 rpm for 5 minutes and virus was aliquoted and stored at −80°C. To generate a working stock, virus from the initial electroporation was amplified once on BHK-21 cells, the supernatant was clarified by centrifugation at 1,200 rpm for 5 minutes and virus was aliquoted and stored at −80°C. All work with CHIKV and LACV variants was completed under ABSL3 conditions at the NYU Grossman School of Medicine.

### LACV Sanger sequencing

The genome of all LACV working stocks and mouse and mosquito samples were Sanger sequenced at Genewiz. In brief, viral RNA was extracted with Trizol following the manufacturers’ instructions. cDNA was generated using the Maxima H First Strand cDNA kit (Thermo) and the S, M, and L segments were amplified by Phusion PCR (Thermo) using the primers in **Table 2**. PCR products were visualized on a 0.8% agarose gel, PCR-column purified (Macherey-Nagel), and sent for Sanger sequencing at Genewiz with the primers in **Table 2**.

**Table 2:**
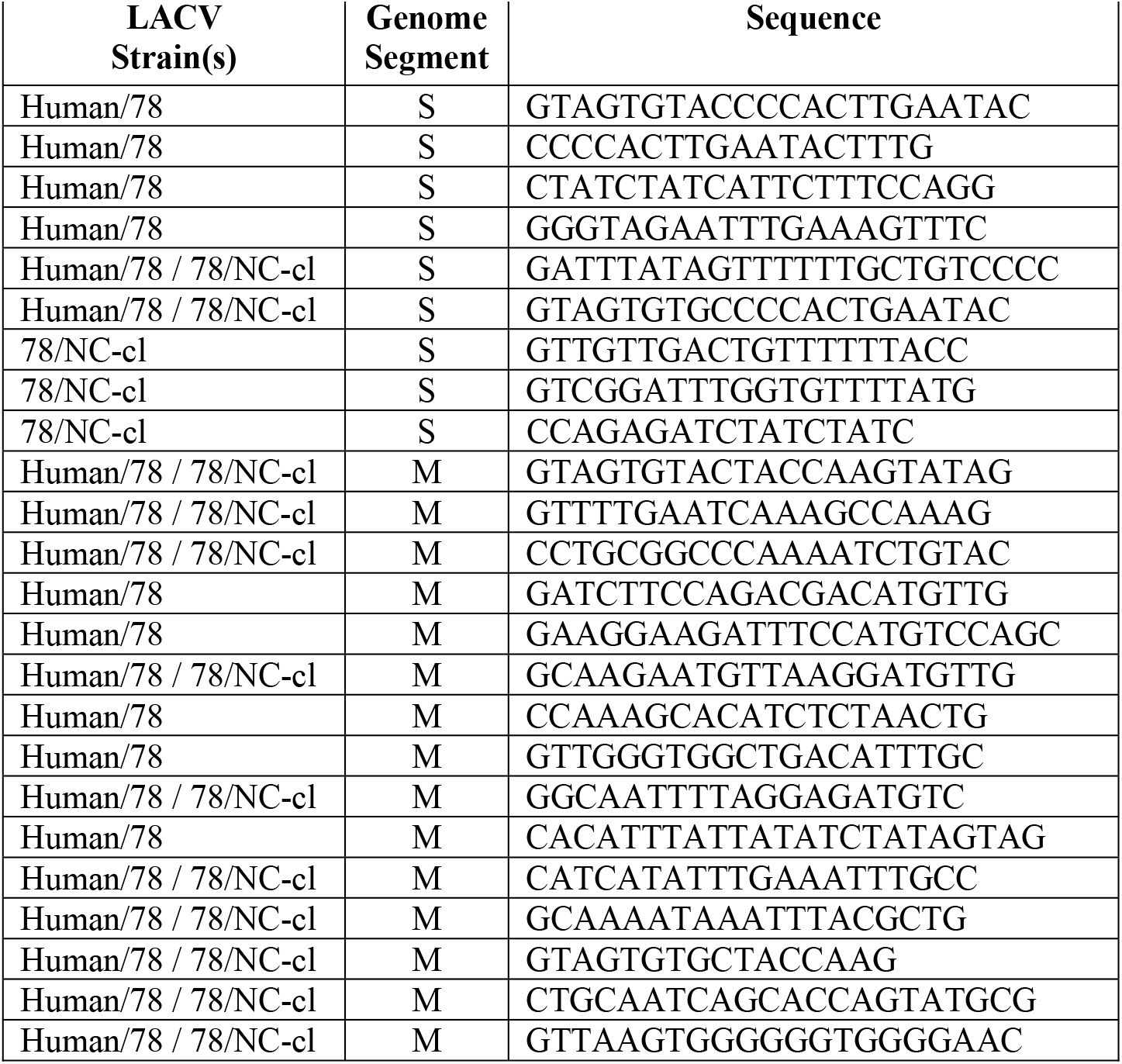
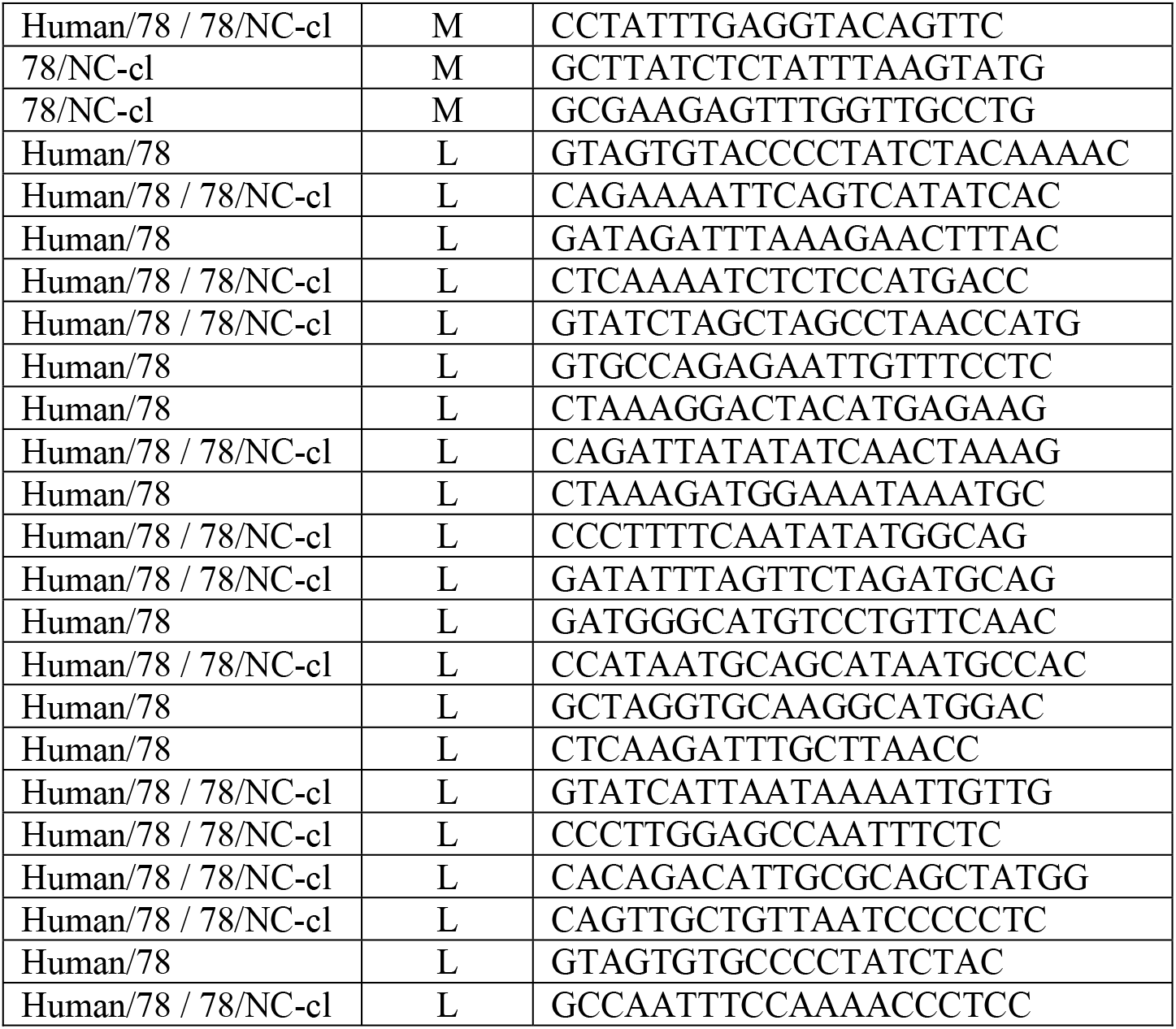
Sequencing primers used in this study.

### Plaque assay

Infectious virus was quantified using a plaque assay where 10-fold serial dilutions of each sample with the virus in DMEM were added to a monolayer of Vero cells. After incubating the virus and cells for 1 hour at 37°C, a media containing 0.8% agarose and DMEM with 2% FBS was added for 3 days at 37°C. The cells were then fixed with 4% formalin for 1 hour, after which the agarose was removed and the plaques were visualized using crystal violet. Viral titers were determined using the lowest countable dilution.

### Cholesterol depletion and complementation assays

BHK-21 cells (20,000 cells/well) were seeded in a Costar 96-well plates and incubated for 24 hours at 37°C. The cells were pretreated with increasing concentrations of methyl-β-cyclodextrin (MβCD) for 1 hour at 37°C. Following incubation, cells were washed once with phosphate buffered saline (PBS), and then incubated with each virus at an MOI = 0.1 for 1 hour at 37°C. After incubation, media containing ammonium chloride was added to a final concentration of 20 mM and the cells were incubated at 37°C for 24 hours. The cells were then fixed with 4% paraformaldehyde (PFA), stained with 4’,6-diamidino-2-phenylindole (DAPI, Thermo Scientific) and LACV antibody as described below, and quantified using a Cell-Insight CX7 High-content microscope (Thermo-Scientific). Cholesterol complementation assay was completed as described above, with the additional step of adding 200 μg/mL of water-soluble cholesterol (Sigma) for 1 hour at 37°C before infection.

### Intracellular LACV staining

Cells were fixed with 4% PFA, incubated with 0.25% TritonX-100 for 10 minutes followed by blocking buffer (0.2% bovine serum albumin, 0.05% saponin) for 1 hour. Cells were then incubated with LACV antisera (a gift from Dr. Karin Peterson, NIH) diluted in blocking buffer for 2 hours. After three PBS washes, cells were then stained with a goat anti-rabbit IgG-Alexa488 secondary antibody (Invitrogen) and DAPI for 1 hour at room temperature.

### Delipidated insect cell assay

Delipidated C6/36 cells were generated as previously described (21). In brief, FBS was delipidated by continuous mixing with 3% (w/v) Cab-O-Sil (ACROS Organics) at 4°C for 48 hours. The FBS was sterile filtered and C6/36 cells were passaged four times in the presence of delipidated serum. C6/36 insect cells were seeded (100,000 cells/well) in a 96-well plate with either complete media or delipidated media and incubated for 24 hours at 28°C. The cells were infected with either LACV WT or CHIKV WT (MOI = 0.1) for 1 hour at 28°C. After washing the cells three times with PBS, either complete media or delipidated media was added back to the cells and left incubating at 28°C. At 24 hours post-infection, the supernatants from each well were harvested and infectious titers were determined via plaque assay.

### Drug assays

BHK-21 cells (20,000 cells/well) were pretreated with increasing concentrations of either Imipramine (TCI America), Atorvastatin (TCI America), Lovastatin (TCI America) or DMSO diluted in complete media. After a 1 hour incubation at 37°C, cells were infected with either LACV or CHIKV-ZsGreen diluted in DMEM at an MOI of 0.1 for an additional hour. The cells were then washed with PBS once, and each compound or DMSO in complete media was given to the cells for an additional 24 hour incubation. Following, the cells were fixed with 8% PFA, stained and infected cells were quantified using a Cell-Insight CX7 High-content microscope (Thermo Scientific).

### MTT cell proliferation assay

To measure the cytotoxicity of each compound, BHK-21 cells (20,000 cells/well) were seeded in a 96-well plate in DMEM supplemented with 10% FBS and incubated for 24 hours at 37°C. Cells were then incubated with increasing concentrations of DMSO or each compound for 24 hours at 37°C. An MTT assay was performed following the manufacturer’s guidelines with the CellTiter 96 Non-Radioactive Cell Proliferation Assay (Promega). For the proliferation readout, 15 μl of the dye solution was added to each well and incubated for 1 hour at 37°C. 100 μl of solubilization solution was added to stop the reaction, and absorbance was read at 570 nm using a Perkin Elmer EnVision plate reader.

### LACV growth curves

BHK-21 (50,000 cells/well in a 24-well plate) were infected with wild-type LACV or each Gc *ij* loop variant at an MOI = 0.1 for 1 hour followed by two washes with PBS and complete media added. Samples of the supernatant were collected at various time points and viral titers were quantified by plaque assay as described above.

### Mouse Infections

Individual C57BL/6J (Jackson Laboratory) breeding pairs were set up to use each liter as a biological replicate. Within one liter, male and female 7-8 day old mice were infected with 50 PFU of wild-type LACV or each *ij* loop variant subcutaneously in the back. Each virus was represented at least one within each liter. Mice were monitored daily then euthanized 3 days post-infection. The brain was harvested, collected in DMEM with 2% serum and homogenized using a Tissue-Lyser II (QIAGEN). Samples were centrifuged at 8,000 rpm for 8 minutes and viral titers were quantified using a plaque assay as described above. All animal experiments were completed in accordance with the NYU Grossman School of Medicine Institutional Animal Care and Use Committee (IACUC) guidelines (protocol no. IA16-01783).

### Mosquito Infections

*Aedes albopictus* (New York City, USA) and *Aedes aegypti* (Poza Rica, Mexico) mosquitoes were previously described (38) (28) (48). In brief, female mosquitoes were fed each virus (10^6^ PFU/mL) diluted in PBS-washed sheep blood containing 5 mM ATP. Mosquitoes were fed for 30 minutes, anesthetized at 4°C, and engorged females sorted into cups. Mosquitoes were incubated at 28°C with 70% relative humidity and 12h:12h diurnal light cycle with 10% sucrose applied ad libitum. At 7 days post-infection, mosquitoes were cold anesthetized and legs and wings removed. The bodies and legs and wings were placed in 200μl PBS, and both were ground in a Tissue-Lyser II and clarified by centrifugation. Viral titers were quantified by plaque assay as described above.

### Protein structures and sequence alignments

The CHIKV E1 glycoprotein (PDB: 3N42), LACV Gc glycoprotein (PDB: 7A57), and LACV Gc head domain (PDB: 6H3W) were generated in PyMol (Version 2.5.2). The bunyavirus Gc *ij* loop sequence alignment was generated using MegAlign (DNA Star) using the following accession numbers: La Crosse virus (AF528166.1), Bunyamwera virus (NC_001926), California encephalitis virus (NC_055118), Jamestown Canyon virus (U88058), Oropouche virus (KP691604), Schmallenberg virus (NC_043584), Snowshoe hare virus (NC_055196). LACV nucleotide sequencing alignments were generated with SeqMan Ultra (DNA Star).

### Statistics and data analysis

All data were analyzed using GraphPad Prism (Version 9.3.1). All *in vitro* experiments were completed with at least two biological replicates and internal technical duplicates or triplicates (exact details can be found in the figure legends). All *in vivo* experiments were completed with at least n=6 for mice and n=10 for mosquito infections. P value of < 0.05 is considered statistically significant.

